# Photocrosslinkable silk fibroin-hyaluronic acid hybrid hydrogels enable chondrocyte-driven matrix deposition and mechanical maturation for cartilage tissue engineering

**DOI:** 10.64898/2026.04.13.718180

**Authors:** Forough Rasoulian, Pejman Ghaffari-Bohlouli, Alexander Otahal, Christoph Bauer, Mohaddeseh Shahabi Nejad, Martin Klein, Armin Shavandi, Abolfazl Heydari, Stefan Nehrer

**Affiliations:** Center for Regenerative Medicine, University of Continuing Education Krems, 3500, Krems, Austria; Université Libre de Bruxelles (ULB), École Polytechnique de Bruxelles, 3BIO-BioMatter, Avenue F.D. Roosevelt, 50-CP 165/61, B-1050 Brussels, Belgium; Polymer Institute of the Slovak Academy of Sciences, Dúbravská cesta 9, 845 41, Bratislava, Slovakia; Institute of Histology and Embryology, Faculty of Medicine, Comenius University in Bratislava, Špitálska Street 24, 813 72 Bratislava, Slovakia; Central European Institute of Technology, Brno University of Technology, 612 00 Brno, Czech Republic

**Keywords:** Photocrosslinkable hydrogel, Silk fibroin, Hyaluronic acid, Articular cartilage regeneration, Chondrocytes, Regenerative medicine

## Abstract

Articular cartilage has limited self-repair capacity, and current treatments fail to fully restore its structure and function. 3D hydrogels that support chondrocyte viability and extracellular matrix (ECM) deposition offer a promising strategy for cartilage regeneration. Here, we developed a photo-crosslinkable silk fibroin–hyaluronic acid hydrogel for 3D encapsulation of primary human chondrocytes. Hydrogels were formulated with varying silk fibroin methacrylate (SilMA, 10-20% w/v) and hyaluronic acid methacrylate (HAMA, 1-2% w/v) concentrations and characterized for rheological, mechanical, and morphological properties. All SilMA-HAMA hydrogel formulations exhibited shear-thinning behavior and rapidly gelled (<20 s) under UV irradiation while maintaining high porosity, thereby ensuring injectability and efficient nutrient diffusion. Notably, the Young’s modulus of the cell-laden scaffolds increased from ∼18 kPa to ∼1200 kPa over culture, indicating mechanical maturation driven by chondrocyte-mediated matrix deposition. This maturation was further confirmed by histological analysis and qPCR, which demonstrated enhanced ECM production and chondrogenic gene expression. Taken together, these results highlight SilMA-HAMA hydrogels as a promising biomimetic platform that couples mechanical reinforcement with biological functionality for cartilage tissue engineering.

## 1. Introduction

Tissue engineering of cartilage using cell-based scaffold strategies has emerged as a promising approach to address the limited regenerative capacity of damaged articular cartilage (1, 2). Owing to its avascular nature, low cellularity, and restricted chondrocyte proliferation, articular cartilage exhibits minimal intrinsic self-repair capability, frequently resulting in progressive cartilage deterioration, subchondral bone remodeling, and eventual development of osteoarthritis (OA) (3). OA affects more than 250 million individuals worldwide (4), and its incidence is projected to rise due to aging populations and increasing obesity rates, posing a significant clinical and socioeconomic burden (5). Current clinical interventions, including microfracture, osteochondral autograft or allograft transplantation, and autologous chondrocyte implantation, are primarily aimed at symptom relief and defect filling rather than true restoration of native hyaline cartilage (5, 6). These approaches are often limited by donor-site morbidity, limited tissue availability, fibrocartilage formation, immune-related complications, and variable long-term outcomes (7, 8). Consequently, cartilage tissue engineering seeks to regenerate functional cartilage by developing biomimetic scaffolds capable of maintaining the chondrocyte phenotype, supporting cell proliferation, and promoting deposition of cartilage-specific extracellular matrix (ECM) components, particularly collagen type II and aggrecan (9). For successful functional regeneration, such scaffolds must combine biocompatibility with mechanical properties that withstand physiological loading while guiding tissue maturation.

Silk fibroin (SF), a natural protein derived from *Bombyx mori* silkworms, has gained significant attention in cartilage tissue engineering due to its biocompatibility, biodegradability, and mechanical robustness (10). SF can be processed into hydrogels, sponges, and films that mimic the fibrous architecture of native ECM, supporting chondrocyte proliferation, regulating apoptosis, and promoting ECM production (11, 12). Hyaluronic acid (HA), a major structural component of cartilage ECM, plays a distinct and complementary role by regulating viscoelastic behavior, osmotic swelling pressure, and cell–matrix signaling. Through these functions, HA directly influences chondrocyte phenotype, mechanotransduction, and cartilage-specific ECM deposition (13). The combination of SF and HA has therefore emerged as a promising strategy to integrate the mechanical robustness of SF with the biological functionality of HA. However, conventional SF/HA hydrogels are often formed via covalent network formation enabled by carbodiimide-mediated coupling reactions (14) or aldehyde–amine condensation (15) pathways, in which multifunctional polymer backbones are interconnected through amide or imine linkages. These approaches typically require precise control of reaction conditions during use and may lead to batch-to-batch variability. These limitations highlight the need for a hydrogel precursor that enables rapid, on-demand crosslinking under mild conditions while maintaining cytocompatibility. To address this need, a rapidly photocrosslinkable hybrid hydrogel composed of methacrylated silk fibroin (SilMA) and methacrylated hyaluronic acid (HAMA) was developed, providing a fully formulated system suitable for minimally invasive delivery and controlled in situ gelation.

Hybrid hydrogels of SilMA and HAMA have demonstrated promising properties for tissue engineering. Previous studies showed that SilMA-HAMA hydrogels support fibroblasts and MC3T3 pre-osteoblasts, exhibiting good biocompatibility and cytocompatibility, and rapid photocrosslinking with enhanced mechanical strength (16). They also promoted fibroblast adhesion, proliferation, and morphology, highlighting potential for 3D bioprinting and mechanically tunable scaffolds (17). Importantly, 3D-printed SilMA-HAMA scaffolds functionalized for cartilage repair efficiently recruited endogenous bone marrow-derived mesenchymal stem cells and provided excellent mechanical stability, supporting regenerative outcomes (18). Beyond SilMA-HAMA systems, chemically crosslinked SF/HA hydrogels prepared via EDC/NHS chemistry promoted chondrogenic differentiation of human bone marrow mesenchymal stem cells, increasing expression of GAG, SOX9, COL2, and aggrecan, confirming strong chondrogenic induction (19). Similarly, injectable SF/HA hydrogels crosslinked with glutaraldehyde and sonication demonstrated biocompatibility, sustained anti-inflammatory drug release, and in vivo gel formation, supporting cartilage regeneration (20). Despite these advances, the capacity of SilMA-HAMA hydrogels to encapsulate chondrocytes and promote cartilage-specific extracellular matrix (ECM) deposition has not yet been fully explored.

In this study, we developed an injectable, cell-laden SilMA-HAMA hydrogel specifically designed for cartilage repair. By combining SilMA with HAMA, the hydrogel rapidly photocrosslinks under UV light, allowing precise handling and minimally invasive delivery. Rheological and mechanical analyses were performed to confirm shear-thinning behavior, injectability, and tunable stiffness. Encapsulation of primary human osteoarthritic chondrocytes enabled evaluation of cytocompatibility, metabolic activity, ECM deposition, and chondrogenic gene expression. Inspired by the native composition of articular cartilage, HAMA incorporation enhances viscoelasticity and functional performance (21), while SilMA provides structural support and scaffold complexity (22, 23). Here, we test the hypothesis that matrix deposition by chondrocytes within SilMA-HAMA hydrogels can drive mechanical maturation, a key outcome for functional cartilage tissue engineering. Taken together, these features position SilMA-HAMA hydrogels as a promising biomimetic platform for regenerative cartilage repair.

## 2. Experimental

### 2.1. Materials

Bombyx mori silk cocoons were obtained from Seidentraum Dr. Matias Langer (Baden-Württemberg, Germany). Sodium carbonate (Na_2_CO_3_), lithium bromide (LiBr), glycidyl methacrylate (GMA), methacrylic anhydride (MA), sodium hydroxide (NaOH), ethanol, deuterium oxide (D_2_O), lithium phenyl (2,4,6-trimethylbenzoyl) phosphinate (LAP), the antibiotic mixture (penicillin, streptomycin, Amphotericin B), ascorbic acid, proteinase K, and β-mercaptoethanol were purchased from Sigma-Aldrich. Hyaluronic acid (HA) with a molecular weight (MW) of 30,000 – 50,000 g·mol^−1^ supplied by Anika Therapeutics Inc. (Bedford, MA, USA). Deionized water and phosphate-buffered saline (PBS 1×) were prepared in-house. High-glucose Dulbecco’s Modified Eagle Medium (DMEM), serum-free chondrogenic medium, and fetal calf serum (FCS) were obtained from Gibco, Thermo Fisher Scientific (Germany). Liberase^TM^ was obtained from Roche Diagnostics (Germany). The XTT Cell Proliferation Assay Kit and the Transcriptor First Strand cDNA Synthesis Kit were purchased from Roche (Basel, Switzerland). Live/Dead staining reagents (Calcein-AM and Ethidium Homodimer-1) were obtained from Invitrogen (USA). The Fibrous Tissue Kit was purchased from Qiagen (Hilden, Germany). Cryosectioning medium (Tissue-Tek O.C.T. Compound) was obtained from Sakura Finetek (The Netherlands). Paraformaldehyde (4%) was purchased from Centralchem (Slovakia). Xylene, paraffin, hematoxylin, and eosin staining reagents, Alcian Blue staining reagent, and Toluidine Blue staining reagent were purchased from Merck (Germany). Ethanol (graded series for dehydration and rehydration) was used for histological sample processing. Glass microscope slides were obtained from Thermo Fisher Scientific.

### 2.2. Methacrylation of silk fibroin and hyaluronic acid

#### Methacrylated silk (SilMA)

SilMA was synthesized following a previously reported method (24). Briefly, 40 g of sliced *Bombyx mori* silk cocoons were boiled in 0.05 mol·L^−1^ Na_2_CO_3_ solution at 100 °C for 30 min to remove sericin. The degummed silk was then dried and dissolved in 9.3 mol·L^−1^ LiBr solution at 60 °C for 60 min. GMA was subsequently added to the silk-LiBr solution, and the mixture was stirred at 60 °C for 4 hours. The resulting solution was dialyzed, using dialysis tubing cellulose membrane with a molar mass cutoff of 14 000 g·mol^−1^ (Sigma-Aldrich), against distilled water for 7 days and freeze-dried to yield SilMA, which was stored at 4 °C until further use.

#### Methacrylated hyaluronic acid (HAMA)

HAMA was synthesized following a previously reported method (25). Briefly, HA was dissolved in water at a concentration of 10 mg·L^−1^. MA was then added dropwise to the HA solution at a stoichiometric ratio of 20 : 1 relative to HA monomer units, with constant magnetic stirring in an ice bath. During the addition of MA and throughout the reaction, the pH was maintained at approximately 9 by addition of 5 mol·L^−1^ NaOH solution. The reaction was allowed to proceed for 6 hours at 4 °C. The resulting HAMA was purified by dialysis using dialysis, tubing cellulose membrane with a molar mass cutoff of 14 000 g·mol^−1^ (Sigma-Aldrich), against distilled water and subsequently isolated by lyophilization.

### 2.3. NMR analysis

The ^1^H NMR analysis was conducted by dissolving 5 mg of the polymers in 500 µL of deuterium oxide. The spectra were recorded at 25 °C, using a 400 MHz Varian INOVA spectrometer. The degree of methacrylation of SilMA (*DM*_SilMA_), defined as the percentage of lysine residues in SF functionalized with methacrylate groups, was determined according to the method of Kim et al. (24) by equation 1:

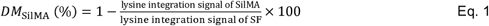

The DM of HAMA (*DM*_HAMA_), defined as the fraction of methacryloyl groups per hyaluronic acid disaccharide repeat unit, was calculated using the method of Bencherif et al. (26) from the ratio of the methacryloyl methyl proton integral (*H*_Me_) to the HA methyl proton integral (*H*_HA_), using equation 2:

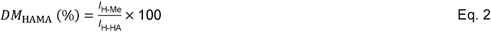

### 2.4. Preparation of cell-free hydrogels

Cell-free SilMA–HAMA hydrogels were prepared according to the compositions summarized in Table 1, in which SilMA concentrations of 10, 15, or 20% (w/v) were combined with HAMA concentrations of 1 or 2% (w/v). As a representative example, for the preparation of the SilMA 15%-HAMA 1% (S15H1) hydrogel, 2 mg of lithium phenyl-2,4,6-trimethylbenzoylphosphinate (LAP) was dissolved in 1 mL of PBS to obtain a 0.2% (w/v) photoinitiator solution, which was protected from light with aluminum foil. Subsequently, 150 mg of SilMA sponge was added to the LAP/PBS solution and stirred at 37 °C until completely dissolved, yielding a homogeneous SilMA precursor solution that was filtered through a 40 µm cell strainer to remove undissolved particles or impurities. Next, 10 mg of HAMA was added to the solution to achieve a final concentration of 1% (w/v) relative to the total volume, and the mixture was vortexed to ensure uniform mixing. Finally, the resulting prepolymer solution was exposed to UV light (365 nm) for 20 s to induce photo-crosslinking, forming the cell-free SilMA-HAMA hydrogel.

**Table 1.** Formulation codes of SilMA-HAMA hydrogel samples, indicating the weight/volume (w/v) concentrations of SilMA and HAMA used in each formulation.

| Sample code | S10H1 | S10H2 | S15H1 | S15H2 | S20H1 | S20H2 |
| --- | --- | --- | --- | --- | --- | --- |
| SiIMA (w/v %) | 10 | 10 | 15 | 15 | 20 | 20 |
| HAMA (w/v %) | 1 | 2 | 1 | 2 | 1 | 2 |

### 2.5. Scanning electron microscopy (SEM)

The SilMA-HAMA hydrogels were observed using a field-emission scanning electron microscope (Hitachi FLEXSEM-1000). Prior to imaging, hydrogel samples were freeze-dried to preserve their internal architecture and subsequently sputter-coated with a thin layer of palladium.

### 2.6. Rheological characterization

Rheological measurements were conducted using an Anton Paar MCR 302 rheometer (Graz, Austria) equipped with a 25 mm parallel-plate geometry. Samples were loaded and conditioned for 2 min prior to measurement, and all experiments were performed at 37 °C using a solvent trap to prevent evaporation. Flow behavior of the hydrogel precursors before UV crosslinking was evaluated by measuring viscosity over a shear rate range of 0.1–1000 s^−1^. Viscoelastic properties of hydrogels were characterized by amplitude sweep tests, during which the storage (G′) and loss (G″) moduli were recorded as a function of shear strain from 0.1% to 1000% at a constant angular frequency of 10 rad·s^−1^; the linear viscoelastic (LVE) region was defined as the strain range in which G′ deviated by less than 5% from its plateau value, and the corresponding yield strain and yield stress were determined. Based on the LVE analysis, a strain amplitude of 1% was selected for frequency sweep measurements, in which G′ and G″ were monitored as a function of angular frequency from 1 to 600 rad·s^−1^ at a constant strain of 1%.

### 2.7. Compression characterization

The compressive mechanical properties of the hydrogels were evaluated by uniaxial compression testing using a Shimadzu AGS-X universal testing machine (Kyoto, Japan) under unconfined conditions. Hydrogel samples were molded into cylindrical specimens with a diameter of 11 mm and a height of 5 mm. Compression tests were performed at a constant crosshead speed of 5 mm·min^−1^ until failure, defined as rupture test, while force–displacement data were continuously recorded and converted into stress–strain curves. Dynamic loading was applied, in which hydrogels were cyclically deformed up to 50% of their initial height (*H*_0_) at a constant speed of 0.1□mm·s^−1^ for 50 consecutive cycles, using S15H1 hydrogel as a representative sample. The compressive Young’s modulus (E) was determined from the linear elastic region of the stress–strain curve, corresponding to 0–5% compressive strain. The slope of this region was obtained by linear regression with a coefficient of determination R^2^ ≥ 0.98. Young’s modulus was calculated using equation 3:

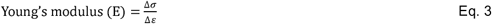

where σ is the compressive stress and ε is the compressive strain.

### 2.8. Zeta Potential

The surface charge of the hydrogels was determined using a SurPASS 3 instrument (Anton Paar GmbH, Austria) equipped with a cylindrical measuring cell, according to the previously reported method (27). Zeta potential measurements were performed using 0.1 mol·L^−1^ NaCl as the electrolyte solution at physiological pH (7.4), under an applied pressure range of 200–600 bar. Prior to measurement, 0.5 g of each hydrogel sample was rinsed three times with 50 mL of 100 mmol·L^−1^ KCl to remove residual salts and to equilibrate the samples with the measurement medium. During analysis, the permeability index of each sample was adjusted to 100 by varying the amount of hydrogel material introduced into the measuring cell. Zeta potential values are reported as the mean ± standard deviation (SD) based on five independent measurements (n = 5).

### 2.9. Water uptake

The water uptake behavior of the samples was evaluated using a modified protocol based on previous reports (28). Briefly, hydrogel samples were first freeze-dried and weighed to obtain their initial dry weight. The dried samples were then immersed in PBS (pH 7.4) at 37 °C for predetermined time intervals of 0.5, 1, 2, 4, 6, 8, 10, and 24 hours. After each interval, the samples were carefully removed, gently blotted with filter paper to remove surface moisture, and weighed to determine the water uptake. The water uptake ratio was calculated using the equation 4:

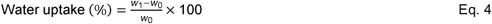

Where *w*_0_ is the dry weight of the hydrogel after freeze-drying, and *w*_1_ is the weight of the hydrogel after water absorption. All measurements were performed in triplicate.

### 2.10. Isolation of chondrocytes from cartilage tissue biopsies

The cartilage tissues were carefully harvested from patients diagnosed with osteoarthritis and undergoing the surgical intervention of knee arthroplasty at the University Hospital Krems. This was done after the necessary ethical clearance, which was registered under the reference number GS1-EK-4/665-2021. The freshly harvested cartilage tissues and adjacent bone pieces were immediately placed into sterile containers filled with PBS. This PBS was supplemented with an antibiotic mixture of penicillin, streptomycin, and Amphotericin B. The samples were then kept at a temperature of 4 °C to maintain their preservation and integrity. To be able to isolate chondrocytes from the surrounding tissue effectively, the tissue was carefully and finely minced into smaller pieces. After this first step, it underwent enzymatic digestion in a nutrient-rich solution containing Liberase. This enzymatic digestion occurred at a controlled temperature of 37 °C and was left overnight with gentle agitation to achieve thorough processing. On the following day, undigested particles that were still in the solution were successfully filtered out by using a 40 μm mesh filter. The isolated cells were then washed, followed by centrifugation, and then they were gently resuspended in a growth medium containing vital antibiotics, namely Amphotericin B, and fetal calf serum (FCS), as well as ascorbic acid. The primary chondrocytes, which were at passage 0, were then seeded into culture flasks at a density of 10,000 cells/cm^2^ of surface area. The culture flasks were maintained in a controlled environment at a temperature of 37 °C within a humidified incubator that had a controlled atmosphere comprising 5% carbon dioxide (CO_2_). In addition, the culture medium was replaced every 2-3 days until the cells had reached a confluence state of about 80%.

### 2.11. Preparation of cell-laden hydrogels

SilMA-HAMA hydrogel precursors were prepared following the same procedure as for the cell-free hydrogels, except that PBS was replaced with serum-free chondrogenic medium. The resulting hydrogel precursor solution was passed through a 40 µm cell strainer (BD, USA) and sterilized by pasteurization technique incubation at 60 °C for 30 min as described in the literature (29). Human chondrocytes were then suspended in the hydrogel precursor at a density of 1 × 10^6^ cells·mL^−1^ and gently mixed to ensure homogeneous cell distribution. Photocrosslinking was performed by exposing the cell-laden precursor to UV light (365 nm, 15 mW/cm^2^) for 20 s. The resulting cell-laden hydrogels were subsequently cultured at 37 °C in a humidified atmosphere containing 5% CO_2_ using high-glucose DMEM supplemented with 10% fetal calf serum (FCS), antibiotics, and 0.05 mg·mL^−1^ascorbic acid.

### 2.12. In vitro cytocompatibility

The metabolic activity of encapsulated human chondrocytes was evaluated using the XTT Cell Proliferation Assay. Cell-laden hydrogels (50 µL) containing human chondrocytes at a density of 1 × 10^6^ cells·mL^−1^ were polymerized directly in 96-well plates. Measurements were performed on days 1, 3, 5, and 7 following the manufacturer’s instructions. Briefly, equal volumes (50 µL each) of XTT reagent and culture medium were mixed and added to each well. After incubation for 5 h at 37 °C in a humidified atmosphere containing 5% CO_2_, absorbance was measured at 492 nm with a reference wavelength of 690 nm using a multimode microplate reader (BioSynergy™ 2, BioTek Instruments, USA). Data were analyzed using Gen5 software, and acellular hydrogels served as negative controls. All measurements were performed in triplicate.

Cell viability was further assessed using a Live/Dead staining assay. For this analysis, 200 µL cell-laden hydrogels containing human chondrocytes at a density of 1 × 10^6^ cells·mL^−1^ were prepared and analyzed on days 0, 5, 7, and 14. The staining solution consisted of 2 µmol·L^−1^ calcein-AM and 4 µmol·L^−1^ ethidium homodimer-1 in PBS, prepared as previously described (29). Prior to staining, samples were gently rinsed with PBS and incubated with 1.5 mL of the staining solution in the dark at room temperature for 35 min. Fluorescence images were acquired using confocal laser scanning microscopy (Leica TCS SP8 MP, Germany).

### 2.13. Sulfated Glycosaminoglycans (sGAG) assay

The sulfated glycosaminoglycan (sGAG) content of chondrocyte-laden SilMA–HAMA hydrogels was quantified following three weeks of culture to assess the chondrogenic capacity of the hydrogels, using the method described by Barbosa et al. (30). The hydrogels were incubated overnight at 56 °C in a solution containing 25 U·mL^−1^ proteinase K to digest both cellular and extracellular components. The enzyme was deactivated by heating the samples to 90 °C for 10 min. After digestion, the mixture was filtered through 0.1□µm ultra-free filter units (Millipore, MA, USA) and centrifuged at 12,000×g for 4 min at room temperature. To determine sGAG levels, 100□µL of the resulting filtrate was combined with 1□mL of 1,9-dimethyl-methylene blue (DMMB) solution to create DMMB-sGAG complexes. These were then pelleted by centrifugation at 12,000×g for 10 min. The pellets were resuspended in a decomplexation buffer and agitated gently for 30 min. Absorbance at 656□nm was measured using an Ultrospec 3300 pro spectrophotometer (Amersham Bioscience, UK). A standard curve based on shark-derived chondroitin sulfate (Sigma-Aldrich, St. Louis, MO, USA) was used for quantification, and all experiments were conducted in triplicate.

### 2.14. RNA isolation

After three weeks of culture, chondrocyte-laden SilMA-HAMA hydrogels were minced into small parts and transferred into tubes with 300□µL of lysis buffer, mixed with 10□µL β-mercaptoethanol and 290□µL RLT buffer from the Fibrous Tissue Kit. To enhance RNA extraction efficiency, the samples were incubated with 20□µL of Proteinase K (included in the same kit) for 30 min, following the manufacturer’s protocol. After elution, purified RNA was stored at −80 °C until reverse transcription for cDNA synthesis.

### 2.15. cDNA synthesis and gene expression analysis

Gene expression was analyzed following the method outlined in a previous study (31). cDNA synthesis was carried out using the Transcriptor First Strand cDNA Synthesis Kit, incorporating RNA carrier (MS2 bacteriophage RNA) to enhance RNA stability. Quantitative real-time PCR (RT-qPCR) was subsequently performed in triplicate using the LightCycler® 96 system (Roche, Basel, Switzerland). The analysis targeted five genes of interest: collagen type II (COL2A1), Aggrecan (ACAN), SRY-Box Transcription Factor 9 (SOX9), collagen type I (COL1A1), and Matrix Metalloproteinase-13 (MMP13). GAPDH (Glyceraldehyde-3-phosphate dehydrogenase) served as the internal reference gene for normalization (Table 2).

**Table 2.**
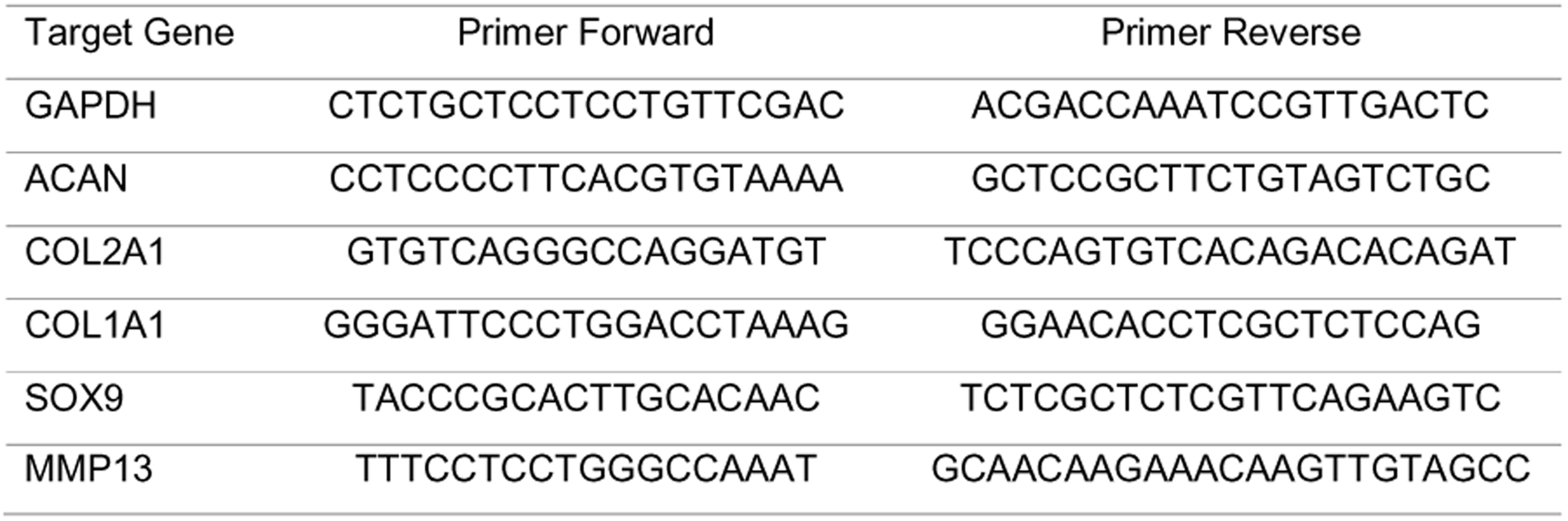
List of primers used in RT-qPCR.

### 2.16. Histology

Hydrogel constructs containing 3D-encapsulated human chondrocytes were harvested on days 1, 7, and 14 for histological evaluation. Samples were gently rinsed with PBS and fixed in 4% paraformaldehyde for 24 h at room temperature. Then the samples were dehydrated through a graded ethanol series up to 100%, cleared with xylene, immersed in paraffin, and embedded in paraffin tissue blocks. Paraffin blocks were sectioned at 5 µm using a rotary microtome. Sections were deparaffinized with Xylene and rehydrated through decreasing ethanol concentrations to 50%, then mounted on glass slides. For each condition and time point, three slides per staining were prepared. General morphology and cell distribution were evaluated using Hematoxylin and Eosin (H&E) staining. Cartilage-like matrix formation was assessed by Alcian Blue staining to detect sGAGs. Toluidine Blue staining was additionally performed to visualize the distribution of proteoglycan-rich ECM. After staining, the slides were mounted with a mounting medium. Images were acquired using a LEICA DM2500 light microscope, and representative photomicrographs were captured using a LEICA DFC290HD digital camera for each condition and time point.

### 2.17. Statistical analysis

GraphPad Prism 10 (GraphPad Software Inc.) was used for statistical analysis. To compare normally distributed datasets, one-way or two-way ANOVA was applied, followed by Tukey’s post hoc test. A p-value of less than 0.05 was regarded as statistically significant. All experiments were performed in triplicate, and the results were reported as mean values ± standard deviation (SD).

## 3. Results and discussion

### 3.1. SilMA and HAMA preparation

SilMA was synthesized by reacting regenerated SF with GMA via epoxide ring-opening of lysine residues (24), as illustrated in Figure 1a. HAMA was prepared by reacting HA with MA (25, 26), as shown in Figure 1b. Successful synthesis of SilMA and HAMA was confirmed by ^1^H NMR spectroscopy. Following GMA addition, SilMA exhibited characteristic vinyl proton peaks at 6.4 and 6.0 ppm and a methyl proton signal at 2.2 ppm, compared to unmodified SF (Figure 1c). HAMA showed vinyl proton peaks at 5.8 and 6.2 ppm and a methyl signal at 2.2 ppm (Figure 1d), confirming methacrylation. These findings were in agreement with previously reported results for SilMA (24) and HAMA (25, 26). The DM value was determined from ^1^H NMR integrals and found to be 40% for SilMA (*DM*_SilMA_) and 40% for HAMA (*DM*_HAMA_).

**Figure 1.**
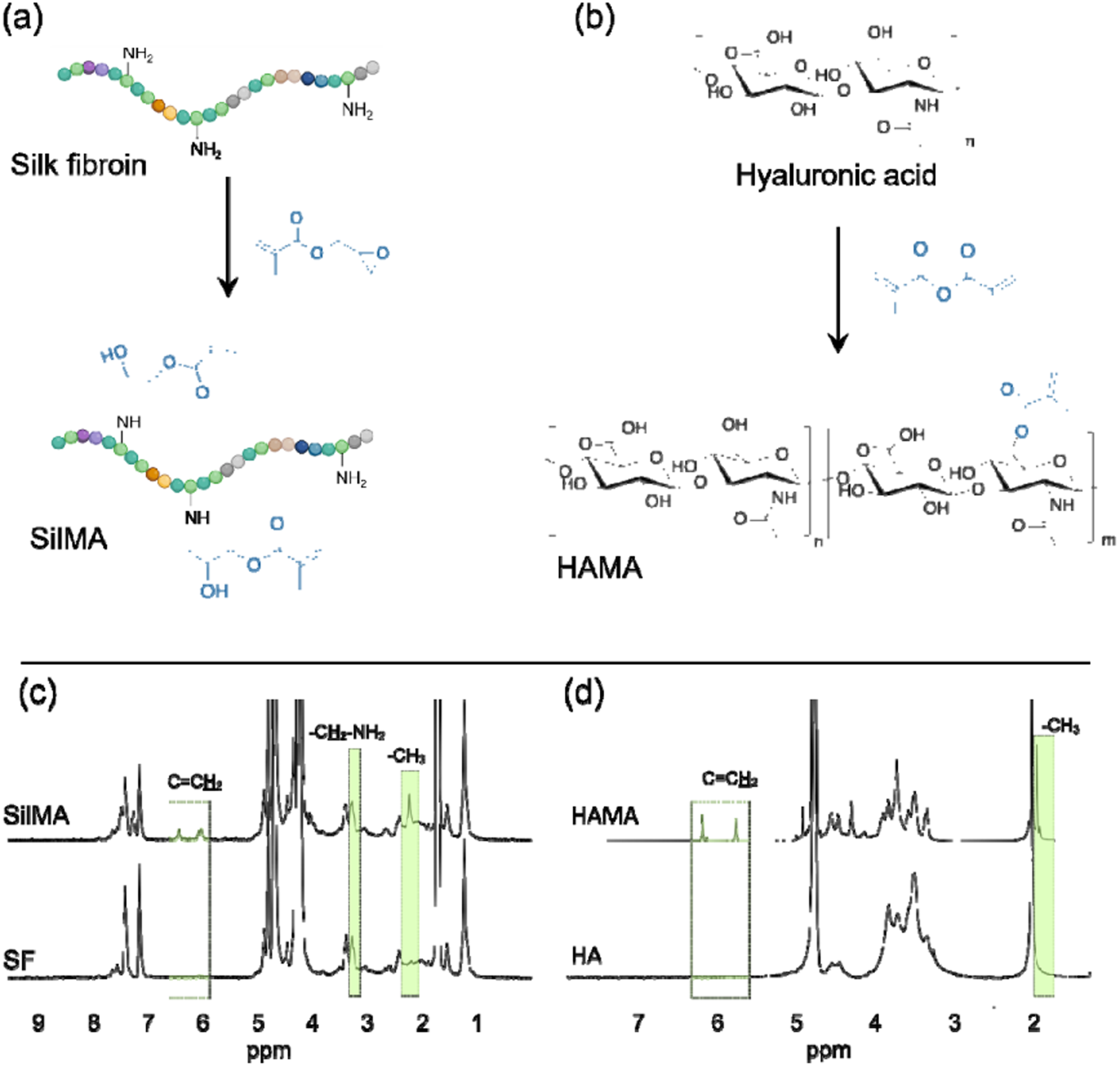
Schematic illustration of the synthesis of (a) methacrylated silk fibroin (SilMA) via reaction of silk fibroin (SF) with glycidyl methacrylate (GMA) and (b) methacrylated hyaluronic acid (HAMA) via reaction of hyaluronic acid (HA) with methacrylic anhydride (MA). ^1^H NMR spectra of (c) SF and SilMA and (d) HA and HAMA recorded in D_2_O.

### 3.2. Cell-free and cell-laden hydrogel preparation and experimental workflow

Figure 2 summarizes the stepwise experimental workflow used to evaluate SilMA-HAMA hydrogels for cartilage tissue engineering. A hydrogel library was generated by varying SilMA (10–20 w/v%) and HAMA (1–2 w/v%) concentrations, yielding six formulations with tunable network density and physicochemical properties. Hydrogel precursor solutions were rapidly crosslinked under 365 nm UV light for 20 s in the presence of a LAP photoinitiator (32). Cell-free hydrogels were initially screened through rheological and compressive testing, microstructural analysis, zeta potential measurements, and water uptake studies to identify formulations with suitable mechanical integrity and hydration behavior. Based on these results and preliminary cytocompatibility assessment via metabolic activity, S15H1 was selected as the optimal formulation and subsequently evaluated in chondrocyte-laden constructs. Biological performance was assessed through cell viability assays, sulfated glycosaminoglycan production, gene expression analysis, histological evaluation, and time-dependent mechanical testing. This rational, multistage approach enables the identification of a SilMA-HAMA hydrogel that combines favorable physicochemical properties with a supportive microenvironment for chondrocyte encapsulation and cartilage regeneration.

**Figure 2.**
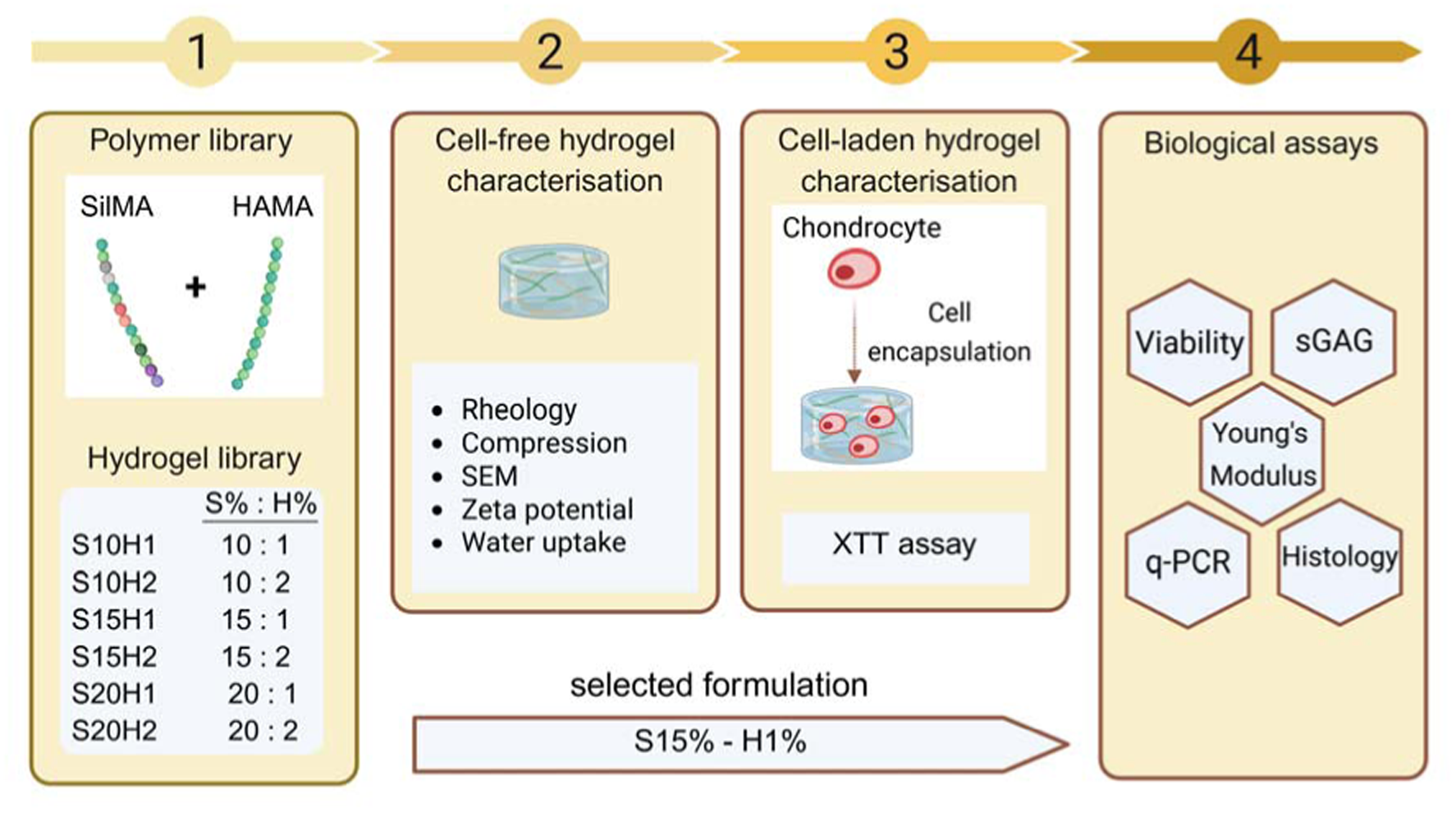
Overview of the experimental workflow for the development and evaluation of SilMA-HAMA hydrogels for chondrocyte encapsulation in cartilage tissue engineering.

### 3.3. Injectability and rheological properties of the hydrogels

The injectability of the hydrogel precursors was first evaluated using a representative formulation (S15H1), which was smoothly extruded through an insulin syringe (Inject-F®, B. Braun, Arolsen, Germany) into PBS solution (Figure 3a) and retained a coherent stream without dispersing, demonstrating both excellent injectability and shape fidelity. This behavior reflects shear-thinning rheology, in which viscosity decreases under applied shear stress and recovers after extrusion, a critical property for minimally invasive hydrogel delivery and regenerating irregular-shaped defects. Rheological measurements confirmed that all SilMA-HAMA precursor formulations exhibited pronounced shear-thinning behavior, as evidenced by a continuous decrease in viscosity with increasing shear rate (Figure 3b). Increasing the SilMA concentration led to a marked rise in precursor viscosity, with the highest values observed for 20% SilMA, reflecting higher polymer content and increased chain entanglement, whereas variations in HAMA concentration (1–2%) had minimal effect on viscosity. These findings indicate that SilMA-HAMA hydrogels possess tunable, shear-thinning properties that enable injectability while maintaining structural integrity prior to photocrosslinking, supporting precise in situ gel formation.

**Figure 3.**
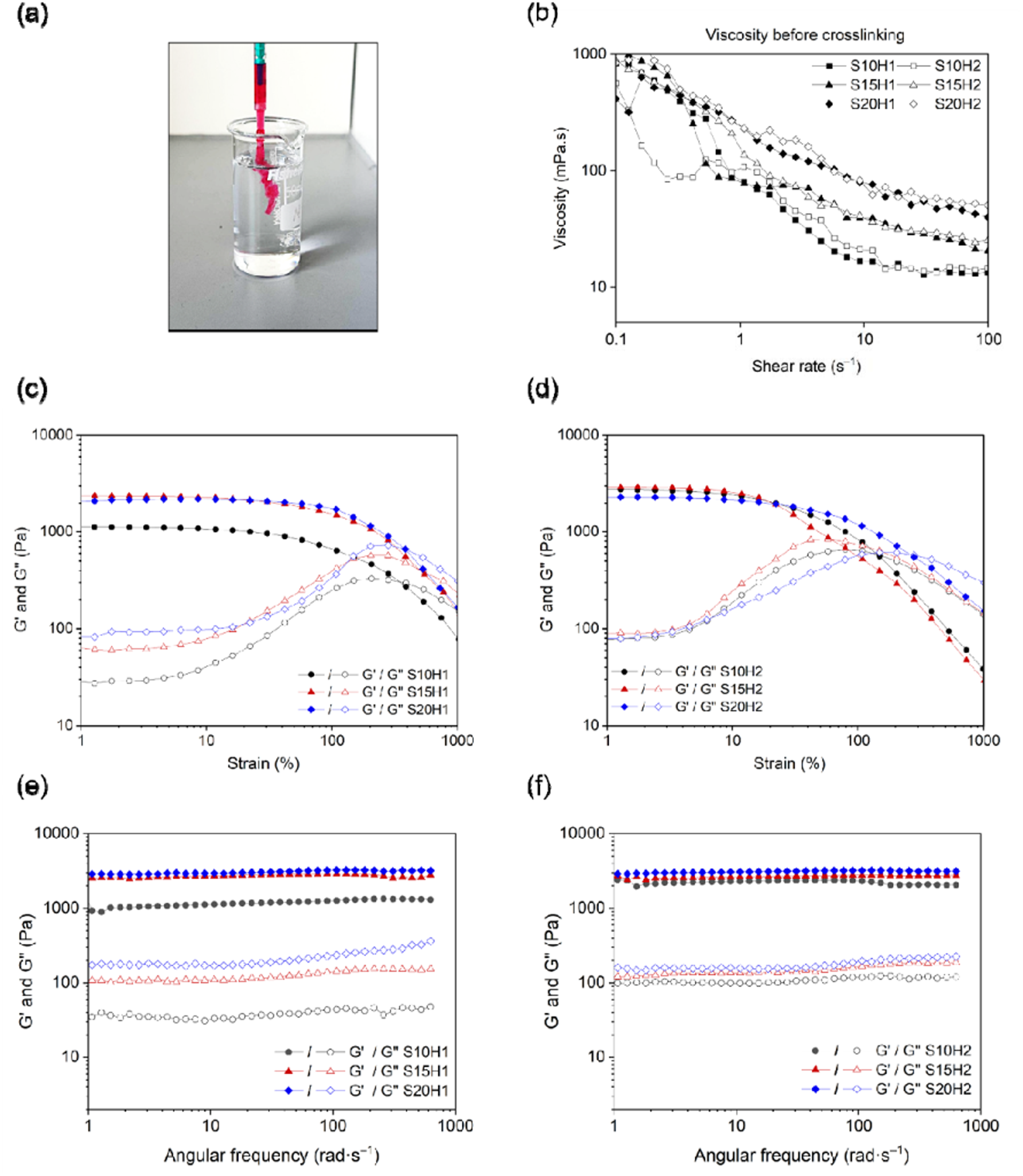
(a) Injectability of SilMA 15%-HAMA 1% (S15H1) hydrogel through a syringe into PBS. (b) Viscosity of SilMA-HAMA hydrogel precursors as a function of shear rate. (c, d) Amplitude sweep analysis of SilMA-HAMA hydrogels at 1 rad·s^−1^ and 37 °C, showing storage modulus (G′) and loss modulus (G″). (e, f) Frequency sweep of SilMA-HAMA hydrogels at 1% strain and 37 °C, demonstrating viscoelastic behavior across a 1–600 rad·s^−1^ range.

Figure 3c-f presents the rheological characteristics of hydrogels. Both amplitude (Figure 3c and d) and frequency sweeps (Figure 3e and f) reveal the gel-like behavior of all samples, since G′ > G″. Amplitude sweep testing was performed to evaluate the viscoelastic behavior and structural integrity of SilMA-HAMA hydrogels, providing insights into their linear viscoelastic (LVE) region, storage modulus (G′), and critical strain. The LVE region, reflecting the strain range over which the hydrogel maintains its network structure, varied substantially across formulations, with S15H1 and S20H1 exhibiting the widest LVE region (∼60%), indicating greater resistance to deformation under small strains, whereas increasing HAMA content in S10H2 and S15H2 reduced the LVE region to ∼10%, suggesting a more brittle network. The storage modulus (G′), representing the elastic stiffness of the hydrogel, increased with both SilMA and HAMA concentrations, with S10H2 and S15H2 showing the highest G′ values (∼2887 Pa), reflecting a stiffer network likely due to increased HAMA-mediated crosslinking, while S10H1 displayed the lowest G′ (1119 Pa). In agreement with our finding, Montaseri et al. reported that incorporating SF in the hydrogel precursor significantly improved its stiffness and elasticity (33). Critical strain, defined at the crossover of G′ and G″ and indicative of the hydrogel tolerance to deformation before yielding, was highest for S15H1 (545%), demonstrating superior network flexibility, whereas HAMA-rich hydrogels (S10H2, S15H2) exhibited lower critical strains (∼150%), consistent with a stiffer, less deformable structure. Furthermore, frequency sweep analysis over a 1–600 rad/s range at 1% strain revealed that G′ and G″ remained largely frequency-independent across all formulations (Figure 3e and f), indicating stable network structures and successful crosslinking through covalent bonding. Collectively, these results suggest that hydrogels with moderate SilMA content and lower HAMA concentrations (e.g., S15H1) achieve an optimal balance between stiffness and extensibility, providing both mechanical support and resilience necessary for cartilage tissue engineering.

### 3.4. Mechanical properties and water uptake of hydrogels

The mechanical properties of SilMA-HAMA hydrogels were evaluated under unconfined compression, with representative compression curves presented in Figure 4a and the quantitative results summarized in Table 3. The hydrogels with different concentrations of SilMA and HAMA were tested, to allow systematic assessment of the influence of polymer composition on mechanical behavior. Young’s modulus, reflecting hydrogel stiffness, clearly depended on polymer composition, increasing from 3 kPa for S10H1 to 64 kPa for S20H2. At a fixed SilMA content, increasing HAMA concentration substantially enhanced stiffness, as seen in S10H1 versus S10H2 (3 kPa vs. 12 kPa), S15H1 versus S15H2 (8 kPa vs. 32 kPa), and S20H1 versus S20H2 (14 kPa vs. 64 kPa), highlighting the pivotal role of HAMA in governing network density and mechanical reinforcement. Compressive stress at break also increased with higher SilMA and HAMA content, ranging from 56 ± 3 kPa (S10H1) to 125 ± 6 kPa (S20H2), whereas compressive strain at break decreased from 80 ± 6% to 49 ± 8%, reflecting the trade-off between stiffness and deformability. Similarly, Wang et al. reported that increasing SF hydrogel concentration enhanced stiffness but slightly reduced flexibility (34), and Phan et al. showed that adding HA to SF hydrogels increased compressive modulus at the expense of elasticity (20), highlighting the trade-off between strength and flexibility. These trends align with the rheological observations: hydrogels with higher HAMA content exhibited higher G′ values and narrower LVE regions, corresponding to stiffer, less extensible networks, while formulations with moderate SilMA and lower HAMA (e.g., S15H1) demonstrated a combination of sufficient stiffness and high deformability, consistent with superior critical strain and wider LVE ranges observed in rheology.

**Figure 4.**
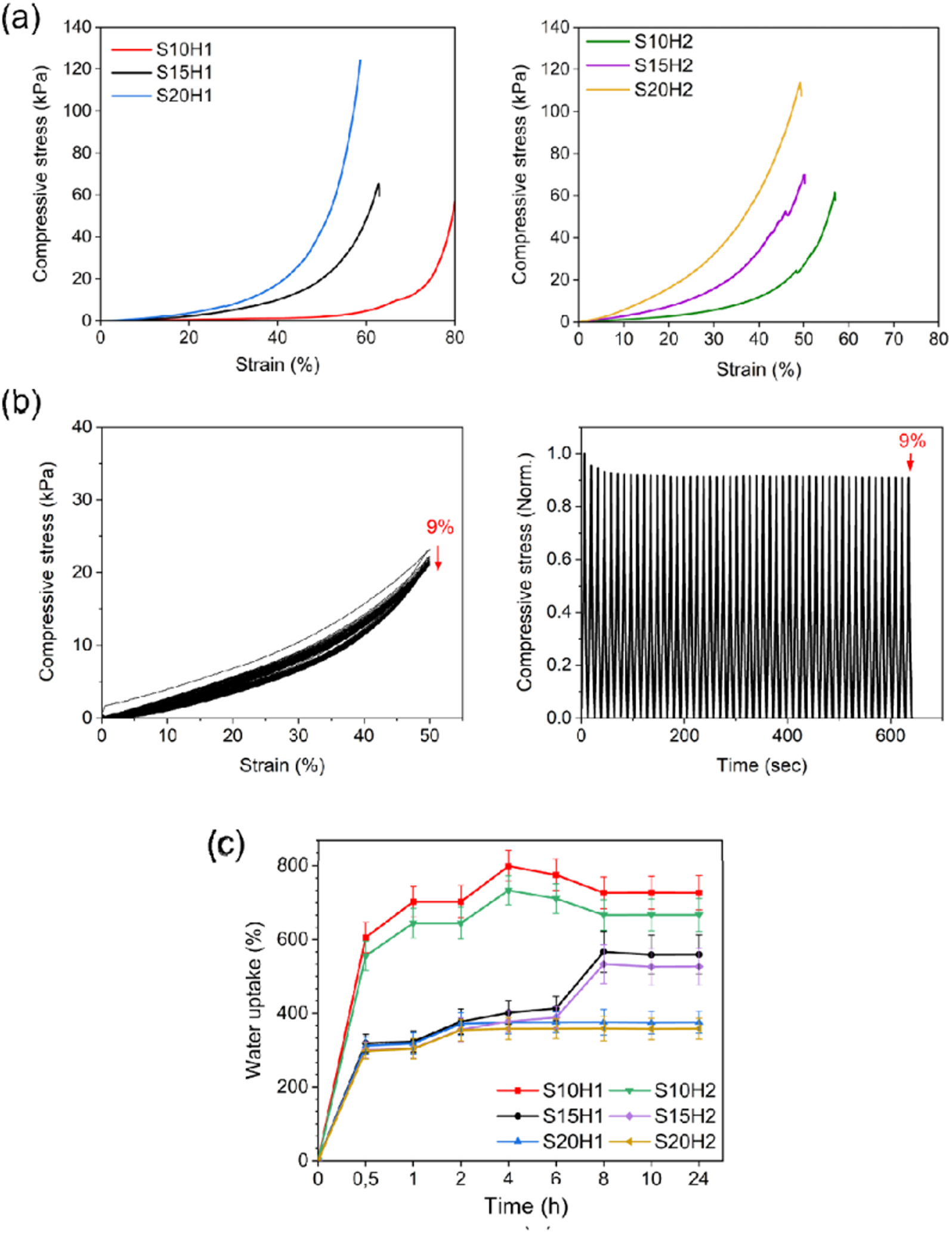
(a) Representative compressive stress–strain curves of cell-free SilMA–HAMA hydrogels obtained from rupture tests. (b) Cyclic compressive deformation behavior of the S15H1 hydrogel, shown as compressive stress–strain curves and compressive stress versus time. (c) Water uptake (in %) of SilMA-HAMA hydrogels.

**Table 3.** Mechanical properties and water uptake of SilMA–HAMA hydrogels with varying polymer compositions.

|  | S10H1 | S10H2 | S15H1 | S15H2 | S20H1 | S20H2 |
| --- | --- | --- | --- | --- | --- | --- |
| Young's modulus (kPa) | $6 \pm 0.4$ | $12 \pm 1.5$ | $8 \pm 1$ | $32 \pm 3.8$ | $14 \pm 1.7$ | $64 \pm 7.5$ |
| Compressive stress at break (kPa) | $56 \pm 3$ | $61 \pm 4$ | $65 \pm 3$ | $70 \pm 6$ | $120 \pm 18$ | $125 \pm 6$ |
| Compressive strain at break (%) | $80 \pm 6$ | $57 \pm 6$ | $63 \pm 8$ | $50 \pm 2$ | $58 \pm 7$ | $49 \pm 8$ |
| Water uptake at 24 h (%) | $727 \pm 46$ | $667 \pm 45$ | $560 \pm 53$ | $527 \pm 50$ | $375 \pm 30$ | $358 \pm 29$ |

Additionally, the resistance of the hydrogel to dynamic compressive loading was evaluated using S15H1. This formulation was selected for detailed characterization based on rheology and compression results. Cyclic compression up to 50% strain for 50 cycles showed a gradual decrease in maximum stress of approximately 9% (Figure 4b), indicating good structural stability under repeated deformation. For comparison, native articular cartilage typically experiences physiological strains of ∼20% in joints (35), suggesting that S15H1 can withstand higher-than-physiological loading while maintaining mechanical integrity.

The water uptake behavior of the SilMA–HAMA hydrogels is presented in Figure 4c. Among the tested formulations, S10H1 and S10H2 exhibited the highest water uptake capacities, with values of 727 ± 46% and 667 ± 45%, respectively. This pronounced swelling can be primarily attributed to the lower overall polymer content of these hybrid hydrogels, which results in a reduced crosslinking density and a more open network structure. As evidenced by the SEM observations and the corresponding mechanical data (Table 3), hydrogels prepared at lower SilMA concentrations displayed larger pore sizes and lower stiffness. Such structural characteristics facilitate greater water penetration and retention within the polymer network, thereby increasing water uptake. In contrast, hydrogels with higher SilMA and/or HAMA contents (S15H1, S15H2, S20H1, and S20H2) exhibited comparable water uptake values, suggesting that beyond a certain polymer concentration threshold, the swelling behavior becomes less sensitive to further increases in crosslinking density. This trend is consistent with previous reports demonstrating an inverse relationship between crosslinking density and swelling ratio in covalently crosslinked hydrogels, where tighter network structures restrict chain mobility and limit water diffusion into the matrix (20, 36). Overall, these results highlight the role of polymer concentration and network architecture in regulating hydrogel hydration, a key parameter influencing nutrient transport and ECM deposition in cell-laden systems intended for cartilage tissue engineering.

### 3.5. Morphology of hydrogels

Scaffold morphology is an important determinant in cartilage tissue engineering, as pore size, porosity, and interconnectivity directly regulate nutrient diffusion, waste transport, cell adhesion, and ECM deposition (37). SEM images (Figure 5) revealed that all SilMA-HAMA hydrogel formulations exhibited a well-defined, highly interconnected porous architecture. Increasing the SilMA concentration from 10% to 20% resulted in a denser polymer network, likely due to enhanced crosslink density associated with SF β-sheet formation, leading to reduced pore size and overall porosity, in agreement with the previous findings (38). In contrast, variations in HAMA concentration (1–2%) had minimal influence on pore morphology, suggesting that the relatively low HAMA content did not significantly affect network formation.

**Figure 5.**
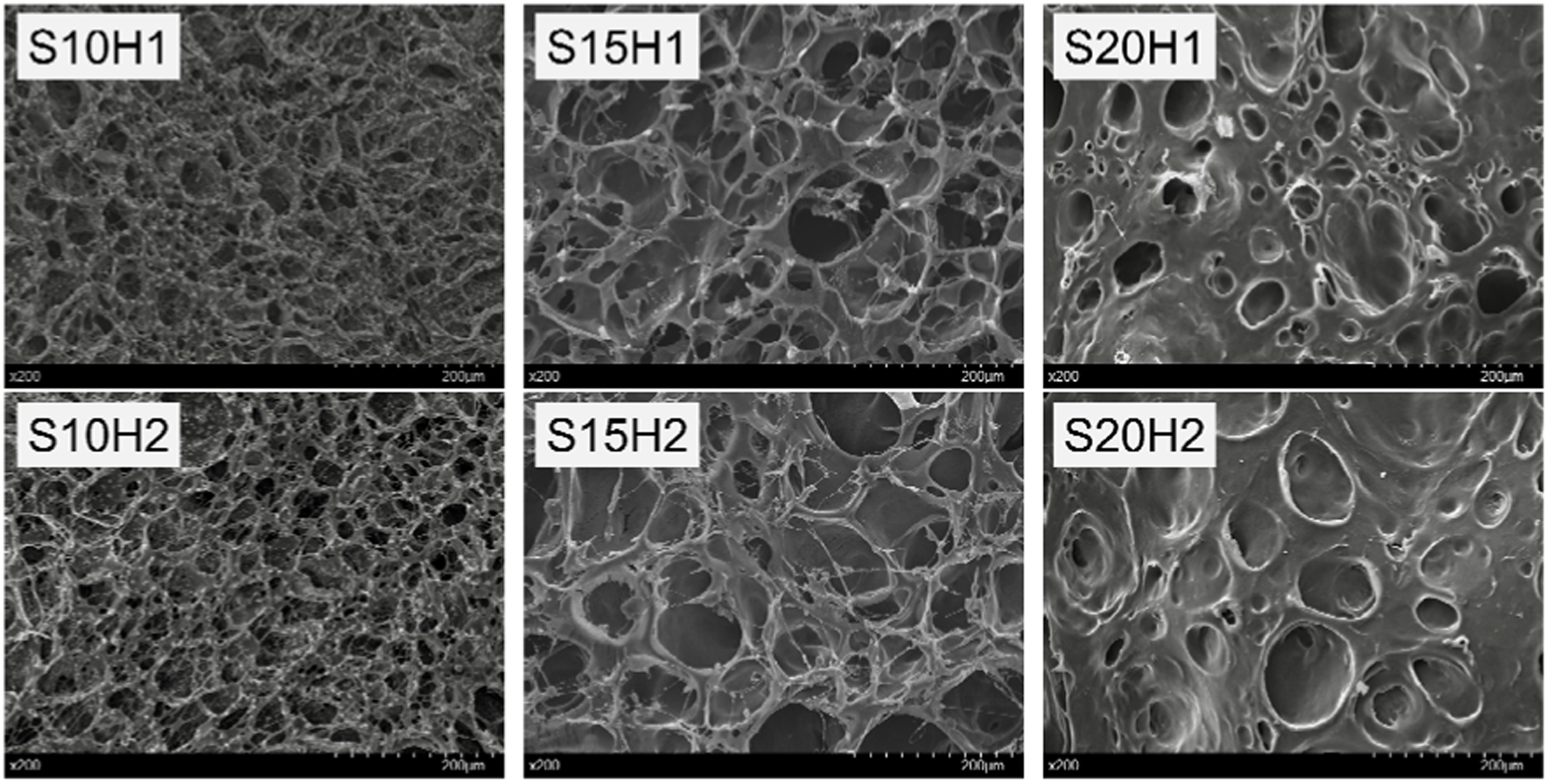
SEM images of SilMA-HAMA hydrogels. Each image includes a 200 μm scale bar.

### 3.6. Zeta potential of SilMA–HAMA hydrogels

The surface zeta potential of SilMA–HAMA hydrogels (S15H1, S15H2, S20H2) was measured, in 0.1 mol·L^−1^ NaCl as the background electrolyte to approximate physiological ionic strength and reduce artifacts due to hypoosmotic conditions (27). SilMA 15% hydrogel, measured as a reference, exhibited a slightly negative zeta potential of approximately −4 ± 1 mV, indicating a weakly anionic surface under the tested conditions. This value is markedly lower than the zeta potential reported for SilMA dispersed in buffer (approximately −23 to −27 mV), suggesting that hydrogel network formation and polymer immobilization significantly reduce the apparent surface charge compared to non-gelled SilMA systems (39). Upon incorporation of HAMA, the surface charge of the hydrogels became more negative. Specifically, SilMA 15% containing 1% HAMA (S15H1) and 2% HAMA (S15H2) both showed a zeta potential of approximately −10 ± 2 mV, demonstrating an increase in negative surface charge compared to SilMA alone. This shift is consistent with the inherently anionic nature of HA, as HA-based hydrogels have been reported to possess negative zeta potentials (40). When the SilMA concentration was raised to 20% with 2% HAMA (S20H2), the surface potential shifted to a less negative value (−5 ± 1 mV). This attenuation of surface negativity is likely attributable to increased SilMA network density and crosslinking, which can shield charged moieties and reduce their effective exposure at the hydrogel–solution interface.

The observed trends suggest that HAMA incorporation effectively increases hydrogel anionicity, a feature that may influence cell–material interactions during cartilage tissue engineering. Negatively charged hydrogel surfaces can mimic aspects of the native ECM, where glycosaminoglycans (GAGs) such as HA contribute to an overall anionic microenvironment (41, 42).

### 3.7. Metabolic activity and cell viability

The metabolic activity of 3D-encapsulated chondrocytes within SilMA-HAMA hydrogels was investigated via the XTT assay. Cell-laden hydrogels were cultured for 7 days, during which metabolic activity was monitored at days 1, 3, 5, and 7. In parallel, cytocompatibility was evaluated through Live/Dead staining, providing a complementary view of cell health and functional activity over time.

An XTT assay was conducted to determine the metabolic activity of chondrocyte-laden SilMA (10%, 15%, and 20%) and HAMA (1% and 2%) (Figure 6a). In the SilMA 10%-HAMA 1% (S10H1) and SilMA 15%-HAMA 1% (S15H1) formulations, in which cellular metabolic activity was found to be increased, the results showed a similar level of metabolic activity for 7 days. Statistical analysis confirmed significant differences between the different formulations of SilMA-HAMA hydrogels at different time points, thus proving that the composition of the hydrogels affected metabolic activity. On the other hand, S20H1 and 2 had relatively low metabolic activity suggesting that a higher SilMA concentration could negatively impact cellular activity. During the 3 to 7-day period, S10H 1/2 and S15H1 hydrogels showed progressive growth in chondrocyte metabolic activity. The values of absorbance measured in the XTT assay (Figure 6a) indicate that the combinations of hydrogels support cellular proliferation. Aqueous SF solutions are generally prepared from concentrated salt solutions of LiBr, and lithium thiocyanate (LiSCN), and the latter can apply negative pressure on cells in chemical modification procedures (43). Subsequently, exposure to such hypertonic conditions for an extended duration could harm cell viability. Earlier studies have shown that residual GMA causes elevated cell death in fibroblast cells (44). Therefore, we optimized the dialysis period of SilMA to eliminate residual GMA and reduce any harmful effects. Determining the right concentration of SilMA and applying suitable purification techniques can improve hydrogel cytocompatibility.

**Figure 6.**
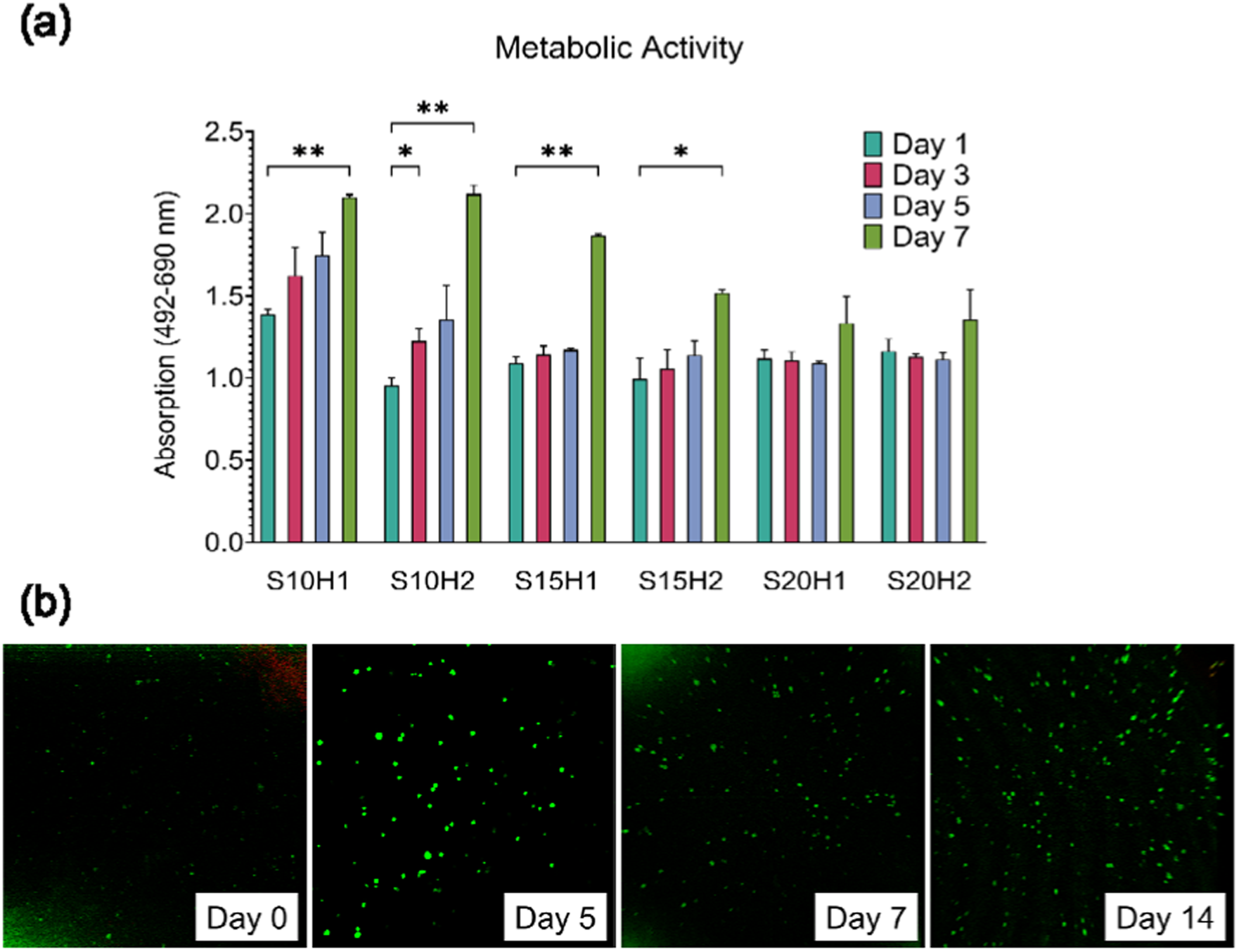
(a) XTT assay showing metabolic activity of human chondrocytes over 1, 3, 5, and 7 days in SilMA- HAMA hydrogels (mean ± SD, *p ≤ 0.05, **p ≤ 0.005). (b) Confocal Live/Dead staining reveals proliferation of chondrocytes in SilMA 15%–HAMA 1% (S15H1) after 7 and 14 days. Scale bar = 200 µm.

Cellular cytocompatibility of the chondrocyte-laden S15H1 hydrogel formulation was further assessed via Live and Dead staining at Days 0, 5, 7, and 14. Confocal images (Figure 6b) indicated negligible red-colored dead cells and many live cells stained in green with Calcein-AM at all the tested time points, suggesting good cellular viability within the hydrogel matrix. Live/Dead imaging qualitatively demonstrated sustained cell survival throughout the 14-day culture duration, as indicated by an apparent increase in cellular presence in the later intervals. Cell-laden hydrogels exhibited visibly higher cell density at Day 14, thus supporting their suitability for extended culture in cartilage tissue engineering.

### 3.8. Matrix formation

GAGs are the predominant components of the cartilage ECM, supplying hydration, compressive stiffness, and biochemical ligand signalization. To determine whether the chondrocyte-laden S15H1 hydrogel can sustain deposition of the ECM, levels of sGAG content were analyzed at day 0, day 7, and day 21 (Figure 7a). No visible difference in the level of the sGAG was observed when comparing day 0 with day 7, indicating that the cells were getting adjusted to the 3D environment during the first week. However, Significant deposition on day 21 compared with day 7 was observed, indicating massive ECM deposition in the hydrogel. Chondrogenic differentiation of the S15H1 hydrogel-encapsulated chondrocytes was analyzed through measurement of the cartilage-related marker genes (COL2A1, ACAN, SOX9), as well as the fibrosis- and hypertrophy-related markers (MMP13, COL1A1), at day 0, day 7, and day 14 (Figure 7b-f). GAPDH was used as a housekeeping gene. COL2A1, a key protein of the hyaline cartilage ECM, is essential in maintaining chondrogenic activity.

**Figure 7.**
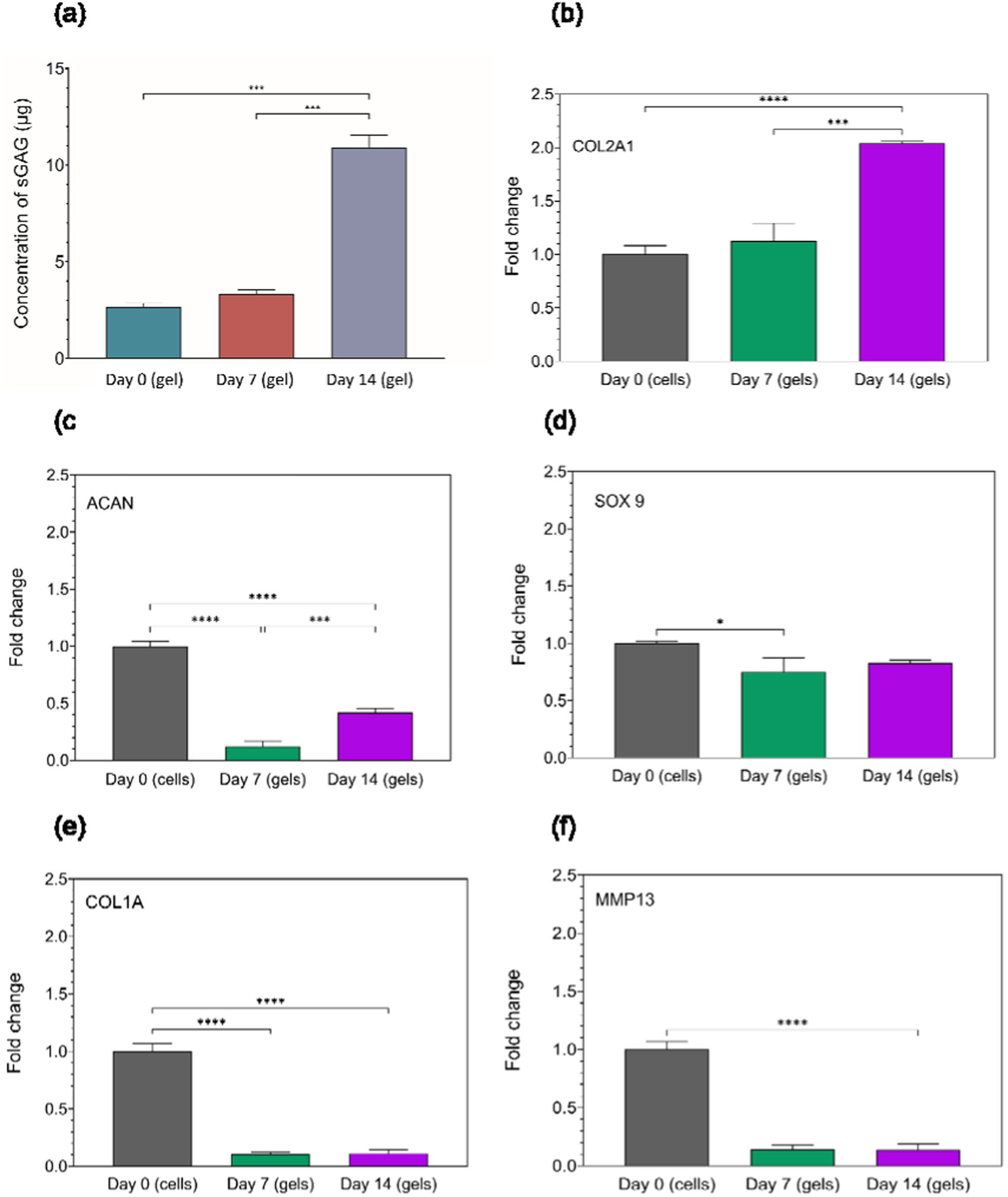
(a) Quantification of sulfated glycosaminoglycan (GAG) content and (b–f) relative gene expression profiles of human chondrocytes encapsulated in methacrylated silk fibroin (SilMA)–methacrylated hyaluronic acid (HAMA) hydrogels during in vitro culture. qPCR analysis shows expression of cartilage-related markers COL2A1 (b), ACAN (c), and SOX9 (d), as well as fibrocartilage-associated COL1A1 (e) and hypertrophy-associated markers MMP13 (f) at indicated time points. Error bars represent the standard deviation. *p ≤ 0.05, **p ≤ 0.01, ***p ≤ 0.001, ****p ≤ 0.0001.

The expression of COL2A1 (Figure 7b) did not change on day 7 compared to 2D culture but significantly increased by day 14. This could suggest that the 3D environment provided by S15H1 hydrogels is more conducive to expressing COL2A1 over time. ACAN (Figure 7c), a critical proteoglycan, contributes to water retention and increases cartilage tissue compressive strength. Despite the significant decrease in ACAN expression on day 7, it indicated recovery by day 14. The reduction at day 7 might indicate an initial adaptation phase, followed by a recovery or adaptation to the 3D environment by day 14. Also, the observed delay in ACAN expression suggests the complexity of chondrocyte maturation and ECM production within the 3D hydrogel environment (45). SOX9 (Figure 7d), a critical transcription factor, significantly affects chondrocyte proliferation and controls cartilage ECM-related genes. SOX9 was detected on day 7 and persisted until day 14, indicating a continuous transcriptional activation of cartilage-specific genes.

COL1A1 is related to fibrocartilage synthesis, while MMP13 is related to cartilage degradation. The expression of both markers (Figure 7e and f) decreased on day 7 and remained low on day 14. This indicates preserving the cartilage phenotype, reducing fibrosis, and preventing degeneration.

Our results show that S15H1 hydrogel promoted ECM production, as supported by the growing amount of sGAG and COL2A1 expression. This finding aligns with the studies on SF and HA-based scaffolds, which provide an environment for sGAG synthesis and ECM deposition over time (46). For example, Li et al. showed that Berberine oleanolic acid-grafted-HA/SF scaffolds promoted ECM deposition under inflammatory conditions, highlighting the importance of adding HA in maintaining the chondrocyte phenotype (47). Our findings are supported by Ziadlou et al., who demonstrated that HA-tyramine/SF composite hydrogels promoted significant chondrogenic differentiation of embedded chondrocytes, with upregulated expression of ACAN, COL2A1, and SOX9 (48). S15H1 hydrogel suppressed the expression of COL1A1 and MMP13, markers of fibrocartilage and matrix degradation. Consistently, Ziadlou et al. found that HA/SF hydrogels supported high COL2A1 and SOX9 expression and moderated COL1A2 levels, promoting hyaline cartilage matrix deposition and reducing fibrocartilage formation. This highlights the role of biomaterial composition in enhancing cartilage matrix production. In the current study, SOX9 expression was maintained, consistent with Ziadlou et al.’s findings, where optimized HA/SF hydrogels supported stable chondrocyte phenotype and ECM deposition over 28 days of culture.

### 3.9. Histology

Histological assessment (Figure 8) of chondrocyte-laden S15H1 hydrogels using H&E, Alcian Blue, and Toluidine Blue demonstrated a coordinated progression of ECM synthesis over the 14-day culture period. H&E staining (Figure 8a) showed that Day 1 hydrogels contained a lightly stained, unorganized matrix with sparse cells. By Day 7, the number of visible nuclei had increased, and regions of denser matrix became apparent, indicating early remodelling. By Day 14, samples displayed pronounced structural organization, including vertically aligned matrix regions and lacuna-like structures, consistent with active chondrocyte matrix interactions. Alcian Blue staining (Figure 8b) revealed a time-dependent increase in acidic GAG deposition. Day 1 constructs showed only faint staining, whereas Day 7 samples exhibited a marked increase in blue coloration distributed throughout the matrix, suggesting early GAG synthesis. By Day 14, Alcian staining became more localized and intense around cells, indicating ECM consolidation and spatial organization of proteoglycans. Toluidine Blue staining (Figure 8c) further confirmed ECM maturation. Day 1 constructs showed minimal staining, while Day 7 samples demonstrated stronger staining around individual cells, indicative of initial sGAG production. Day 14 constructs exhibited the most robust response, with dense pericellular staining and aligned matrix zones, reflecting substantial sulfated proteoglycan deposition and advanced cartilage-like matrix development.

**Figure 8.**
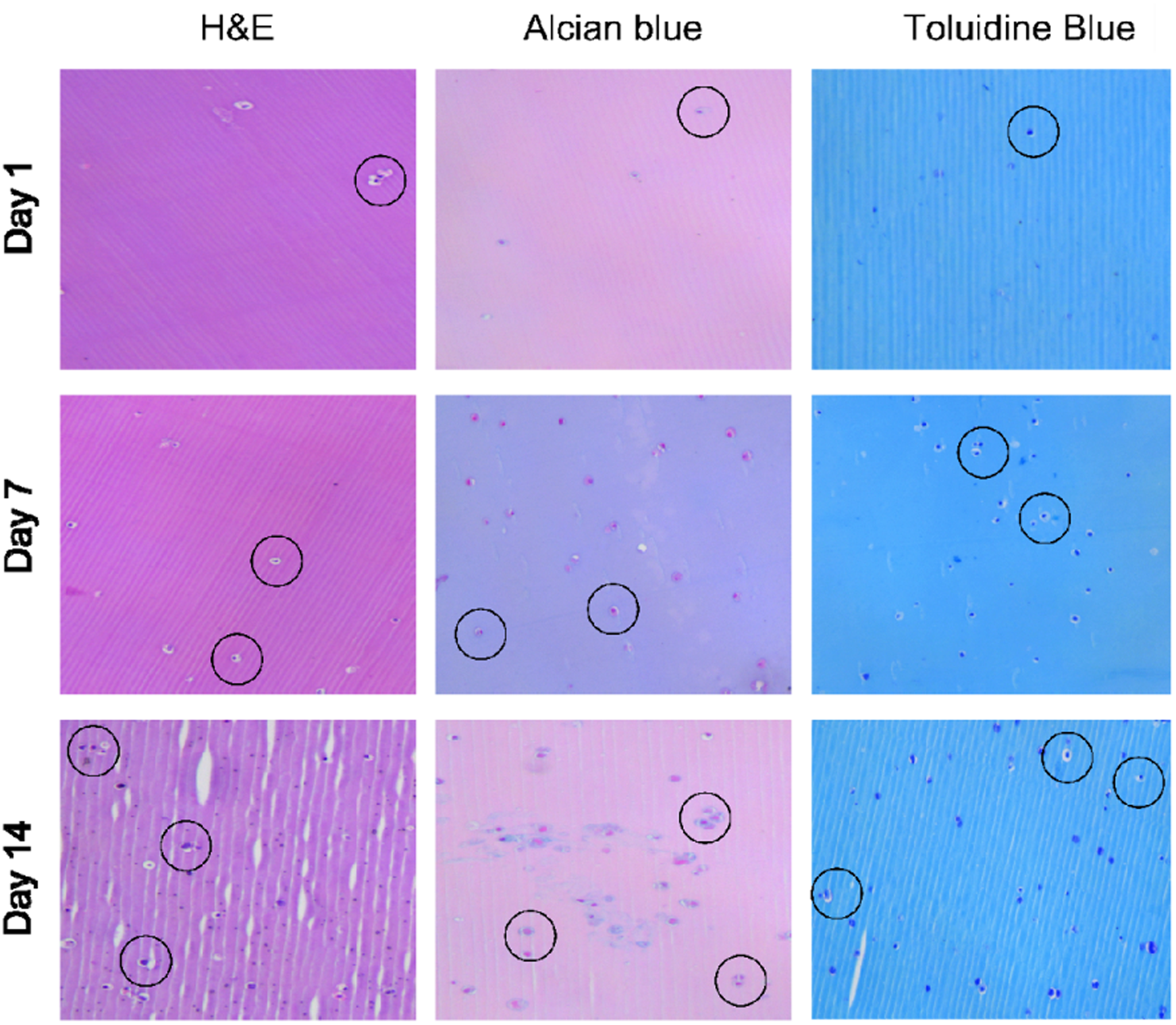
Histological evaluation of chondrocyte-laden S15H1 hydrogels at days 1, 7, and 14. (a) Hematoxylin and Eosin (H&E) staining was performed to assess general morphology and cell distribution, with nuclei stained dark blue/purple and cytoplasm/extracellular components stained pink. (b) Alcian Blue and Nuclear Fast Red staining were used to visualize cartilage-like matrix formation, with nuclei stained red and sulfated glycosaminoglycans (sGAGs) stained blue. (c) Toluidine Blue staining was performed to evaluate the distribution of proteoglycan-rich extracellular matrix. Images were captured at 10× magnification, and scale bars indicate 50 µm.

The combined histological findings demonstrate clear time-dependent ECM development within the constructs. Structural changes observed in H&E staining, such as increased cell density, lacuna-like formation, and emerging matrix alignment, are consistent with early stages of chondrogenic tissue organization described in engineered cartilage systems (49). Biochemical stains further supported this progression. Alcian Blue revealed increasing acidic GAG deposition from Day 1 to Day 14, reflecting the early and intermediate phases of proteoglycan synthesis typical of chondrocyte differentiation (49). The shift from diffuse staining at Day 7 to more localized pericellular accumulation at Day 14 suggests matrix condensation and proteoglycan organization. Toluidine Blue provides more specific information about ECM quality. Its increasing staining intensity over time indicates rising sGAG content, particularly chondroitin sulfate and keratan sulfate, which are essential for the osmotic swelling and mechanical behavior of hyaline cartilage (18). Compared with Alcian Blue (broader acidic GAG detection), Toluidine Blue strongly reflected ECM maturation through increased sulfation. This trend aligns with reports showing that maturing cartilage constructs progressively increase GAG sulfation as they develop toward a more functional phenotype (19). Together, these findings indicate increasing ECM deposition accompanied by improved organization and sulfation hallmarks of early cartilage-like tissue formation.

### 3.10. Mechanical maturation of cell-laden hydrogel

The compressive mechanical properties of the S15H1 hydrogel were evaluated after incubation in culture medium with and without encapsulated chondrocytes to investigate time-dependent mechanical maturation (Figure 9). The freshly prepared hydrogel (day 0) exhibited Young’s modulus values for both cell-free and chondrocyte-laden groups (∼10-20 kPa), with no statistically significant difference between conditions. Under cell-free conditions, a gradual increase in stiffness was observed over time, with Young’s modulus increasing significantly to approximately 360 kPa after 7 days and further to ∼430 kPa by day 14 (compared to day 0). This progressive stiffening in the absence of cells is consistent with the well-documented behavior of silk fibroin-based hydrogels, which undergo gradual secondary structural rearrangement characterized by increased β-sheet content during incubation in aqueous media. Such β-sheet formation enhances crosslinking within the network, leading to increased mechanical stability and stiffness over time (34).

**Figure 9.**
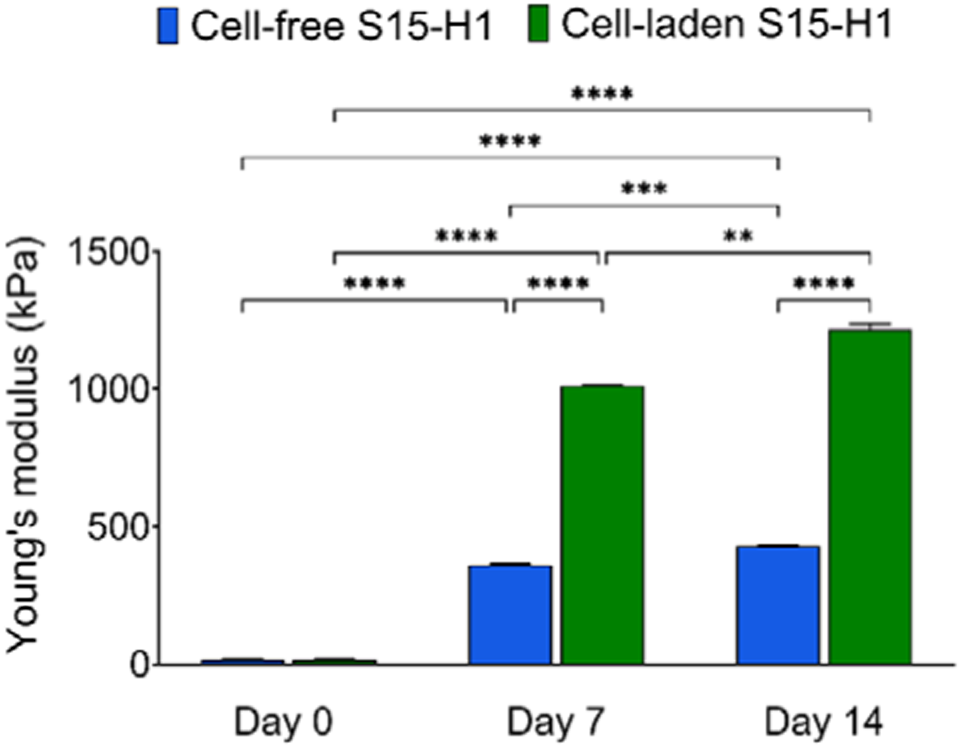
Young’s modulus (kPa) of S15H1 hydrogels under cell-free and chondrocyte-laden conditions after 0, 7, and 14 days of incubation in culture medium. Data are presented as mean ± SD (n = 3). Statistical significance was determined at p < 0.05.

In the presence of chondrocytes, mechanical reinforcement was markedly amplified (Figure 9). Chondrocyte-laden S15H1 hydrogels exhibited a significantly higher Young’s modulus compared to cell-free controls at both day 7 and day 14 (p < 0.05), reaching approximately 1010 kPa after 7 days and further increasing to ∼1200 kPa by day 14.. This corresponds to nearly a threefold increase relative to the corresponding cell-free hydrogels at each time point.. This pronounced mechanical maturation is primarily attributed to extracellular matrix (ECM) deposition by encapsulated chondrocytes, including collagen type II and sGAGs, which integrate within the hydrogel network and reinforce its structure. Notably, the observed Young’s modulus approaches that of native articular cartilage (1.03 ± 0.48 MPa) (50), indicating that the hydrogels provide a mechanically relevant environment for cartilage regeneration. These findings are consistent with previous reports by Wan et al., who found that chondrocyte-laden chitosan/SF hydrogels exhibited enhanced mechanical strength over time due to increased ECM deposition and collagen production (51). Collectively, these results demonstrate that the combination of intrinsic SF network stiffening and cell-mediated ECM deposition enables progressive mechanical maturation, supporting the potential of S15H1 hydrogels as a scaffold for cartilage tissue engineering.

## 4. Conclusions

In this study, we developed and systematically evaluated a photocrosslinkable silk fibroin methacrylate–hyaluronic acid methacrylate (SilMA-HAMA) hydrogel as a biomimetic scaffold for cartilage tissue engineering. By modulating the relative concentrations of SilMA and HAMA, a hydrogel library with tunable network architecture and mechanical properties was established. All formulations exhibited rapid UV-induced crosslinking and a highly porous microstructure, supporting mass transport and nutrient diffusion within three-dimensional constructs.

Encapsulation of primary human chondrocytes demonstrated high cytocompatibility, as evidenced by sustained metabolic activity and high cell viability over time. Among the evaluated formulations, S15H1 (SilMA 15% and HAMA 1%) emerged as the optimal composition, providing a favorable balance between mechanical stability and biological performance. Importantly, chondrocyte-laden S15H1 hydrogels displayed pronounced time-dependent mechanical maturation, with Young’s modulus increasing from the kilopascal range to several hundred kilopascals during culture. This progressive stiffening resulted from the combined effects of intrinsic silk fibroin network stabilization and cell-mediated deposition of cartilage-specific extracellular matrix components, including collagen type II and sulfated glycosaminoglycans. Consequently, the mechanical properties of the engineered constructs approached those of native articular cartilage, underscoring their functional relevance.

Overall, these findings demonstrate that SilMA-HAMA hydrogels function as dynamically maturing, cell-instructive scaffolds that effectively integrate physicochemical tunability with robust biological functionality. The injectable and photocrosslinkable nature of this system, together with its capacity to support chondrocyte viability, extracellular matrix formation, and mechanical reinforcement, positions SilMA-HAMA hydrogels as a promising platform for minimally invasive cartilage repair and future translational applications in regenerative medicine.

## Acknowledgment

This work was supported by the fund for the economy and tourism of the state of Lower Austria (WST3-F-5030664/023-2020). This work was also supported by the Slovak Research and Development Agency under the contract number APVV-22-0568, APVV-22-0565, the Slovak Grant Agency VEGA 2/0154/26, and the European Fund for Regional Development under the project number CZ.02.01.01/00/22_008/0004562.

## Ethics statement

Human tissue used in the study was obtained from patients undergoing total knee replacement surgery after written informed consent. A positive ethics vote was obtained from the NÖ Ethikkommission (GS1-EK-4/761-2021).

## Author contributions

Forough Rasoulian: Conceptualization, Investigation, Methodology, Formal analysis, Visualization, writing original draft, Review, and Editing; Pejman Ghaffari-Bohlouli: Methodology ; Alexander Otahal: Supervision, Review and Editing, Validation; Christoph Bauer: Funding acquisition; Armin Shavandi: Review and Editing; Mohaddeseh Shahabi Nejad: Methodology; Martin Klein: Methodology; Abolfazl Heydari: Review and Editing, Validation, Funding acquisition, Stefan Nehrer: Supervision, Funding acquisition, Resources, Review and Editing. All authors critically reviewed the manuscript and agreed on the final version.

## References

1. Zhao X, Hu DA, Wu D, He F, Wang H, Huang L, et al. Applications of Biocompatible Scaffold Materials in Stem Cell-Based Cartilage Tissue Engineering. Front Bioeng Biotechnol. 2021;9:603444.

2. Liang J, Liu P, Yang X, Liu L, Zhang Y, Wang Q, et al. Biomaterial-based scaffolds in promotion of cartilage regeneration: Recent advances and emerging applications. J Orthop Translat. 2023;41:54–62.

3. Pueyo Moliner A, Ito K, Zaucke F, Kelly DJ, de Ruijter M, Malda J. Restoring articular cartilage: insights from structure, composition and development. Nat Rev Rheumatol. 2025;21(5):291–308.

4. Wu R, Guo Y, Chen Y, Zhang J. Osteoarthritis burden and inequality from 1990 to 2021: a systematic analysis for the global burden of disease Study 2021. Sci Rep. 2025;15(1):8305.

5. Scheuing WJ, Reginato AM, Deeb M, Acer Kasman S. The burden of osteoarthritis: Is it a rising problem? Best Pract Res Clin Rheumatol. 2023;37(2):101836.

6. Kutaish H, Klopfenstein A, Obeid Adorisio SN, Tscholl PM, Fucentese S. Current trends in the treatment of focal cartilage lesions: a comprehensive review. EFORT Open Reviews. 2025;10(4):203–12.

7. Laurent A, Abdel-Sayed P, Ducrot A, Hirt-Burri N, Scaletta C, Jaccoud S, et al. Development of Standardized Fetal Progenitor Cell Therapy for Cartilage Regenerative Medicine: Industrial Transposition and Preliminary Safety in Xenogeneic Transplantation. Biomolecules. 2021;11(2):250.

8. Makris EA, Gomoll AH, Malizos KN, Hu JC, Athanasiou KA. Repair and tissue engineering techniques for articular cartilage. Nat Rev Rheumatol. 2015;11(1):21–34.

9. Xiong Z, Hong F, Wu Z, Ren Y, Sun N, Heng BC, et al. Gradient scaffolds for osteochondral tissue engineering and regeneration. Chem Eng J. 2024;498:154797.

10. Zhou Z, Cui J, Wu S, Geng Z, Su J. Silk fibroin-based biomaterials for cartilage/osteochondral repair. Theranostics. 2022;12(11):5103–24.

11. Wu Y, Zhou L, Li Y, Lou X. Osteoblast-derived extracellular matrix coated PLLA/silk fibroin composite nanofibers promote osteogenic differentiation of bone mesenchymal stem cells. J Biomed Mater Res A. 2022;110(3):525–34.

12. Rainbow RS, Kwon H, Foote AT, Preda RC, Kaplan DL, Zeng L. Muscle cell-derived factors inhibit inflammatory stimuli-induced damage in hMSC-derived chondrocytes. Osteoarthritis Cartilage. 2013;21(7):990–8.

13. Wu H, Shen L, Zhu Z, Luo X, Zhai Y, Hua X, et al. A cell-free therapy for articular cartilage repair based on synergistic delivery of SDF-1 & KGN with HA injectable scaffold. Chem Eng J. 2020;393:124649.

14. Yan S, Wang Q, Tariq Z, You R, Li X, Li M, et al. Facile preparation of bioactive silk fibroin/hyaluronic acid hydrogels. Int J Biol Macromol. 2018;118(Pt A):775–82.

15. Zhang Y, Liu L, Guan C, Xu C, Liu Y, Abdullah M, et al. Aloe extract-loaded cationized silk fibroin/oxidized hyaluronic acid injectable self-healing hydrogel for diabetic wound healing. Int J Biol Macromol. 2025;338(Pt 1):149595.

16. Amirian J, Wychowaniec JK, D′este M, Vernengo AJ, Metlova A, Sizovs A, et al. Preparation and Characterization of Photo-Cross-Linkable Methacrylated Silk Fibroin and Methacrylated Hyaluronic Acid Composite Hydrogels. Biomacromolecules. 2024;25(11):7078–97.

17. Li S, Huang C, Liu H, Han X, Wang Z, Huang J, et al. A Silk Fibroin Methacryloyl-Modified Hydrogel Promoting Cell Adhesion for Customized 3D Cell-Laden Structures. Acs Appl Polym Mater. 2022;4(10):7014–24.

18. Shi W, Zhang J, Gao Z, Hu F, Kong S, Hu X, et al. Three-Dimensional Printed Silk Fibroin/Hyaluronic Acid Scaffold with Functionalized Modification Results in Excellent Mechanical Strength and Efficient Endogenous Cell Recruitment for Articular Cartilage Regeneration. Int J Mol Sci. 2024;25(19):10523.

19. Song S, Li B, Gao X, Zhang Z, Zhou Y, Liu X, et al. Bioengineered Silk Fibroin/Hyaluronic Acid Composite Hydrogel for Minimally Invasive Cartilage Repair. Biomed Mater. 2025;20(5):055006.

20. Phan VHG, Murugesan M, Nguyen PPT, Luu CH, Le NH, Nguyen HT, et al. Biomimetic injectable hydrogel based on silk fibroin/hyaluronic acid embedded with methylprednisolone for cartilage regeneration. Colloids and surfaces B, Biointerfaces. 2022;219:112859.

21. Cowman MK, Schmidt TA, Raghavan P, Stecco A. Viscoelastic Properties of Hyaluronan in Physiological Conditions. F1000Res. 2015;4:622.

22. Kim SH, Lee YJ, Chao JR, Kim DY, Sultan MT, Lee HJ, et al. Rapidly photocurable silk fibroin sealant for clinical applications. NPG Asia Materials. 2020;12(1):46.

23. Kim SH, Seo YB, Yeon YK, Lee YJ, Park HS, Sultan MT, et al. 4D-bioprinted silk hydrogels for tissue engineering. Biomaterials. 2020;260:120281.

24. Kim SH, Yeon YK, Lee JM, Chao JR, Lee YJ, Seo YB, et al. Precisely printable and biocompatible silk fibroin bioink for digital light processing 3D printing. Nat Commun. 2018;9(1):1620.

25. Spearman BS, Agrawal NK, Rubiano A, Simmons CS, Mobini S, Schmidt CE. Tunable methacrylated hyaluronic acid-based hydrogels as scaffolds for soft tissue engineering applications. J Biomed Mater Res A. 2020;108(2):279–91.

26. Bencherif SA, Srinivasan A, Horkay F, Hollinger JO, Matyjaszewski K, Washburn NR. Influence of the degree of methacrylation on hyaluronic acid hydrogels properties. Biomaterials. 2008;29(12):1739–49.

27. Dorchei F, Heydari A, Kronekova Z, Kronek J, Pelach M, Cseriova Z, et al. Postmodification with Polycations Enhances Key Properties of Alginate-Based Multicomponent Microcapsules. Biomacromolecules. 2024;25(7):4118–38.

28. Li Y, Zhang J, Wang C, Jiang Z, Lai K, Wang Y, et al. Porous composite hydrogels with improved MSC survival for robust epithelial sealing around implants and M2 macrophage polarization. Acta Biomater. 2023;157:108–23.

29. Kim SH, Hong H, Ajiteru O, Sultan MT, Lee YJ, Lee JS, et al. 3D bioprinted silk fibroin hydrogels for tissue engineering. Nat Protoc. 2021;16(12):5484–532.

30. Barbosa I, Garcia S, Barbier-Chassefière V, Caruelle JP, Martelly I, Papy-García D. Improved and simple micro assay for sulfated glycosaminoglycans quantification in biological extracts and its use in skin and muscle tissue studies. Glycobiology. 2003;13(9):647–53.

31. Bauer C, Niculescu-Morzsa E, Nehrer S. A protocol for gene expression analysis of chondrocytes from bovine osteochondral plugs used for biotribological applications. MethodsX. 2017;4:423–8.

32. Applegate MB, Partlow BP, Coburn J, Marelli B, Pirie C, Pineda R, et al. Photocrosslinking of Silk Fibroin Using Riboflavin for Ocular Prostheses. Adv Mater. 2016;28(12):2417–20.

33. Montaseri Z, Abolmaali SS, Tamaddon AM, Farvadi F. Composite silk fibroin hydrogel scaffolds for cartilage tissue regeneration. J Drug Deliv Sci Technol. 2023;79:104018.

34. Wang T, Li Y, Liu J, Fang Y, Guo W, Liu Y, et al. Intraarticularly injectable silk hydrogel microspheres with enhanced mechanical and structural stability to attenuate osteoarthritis. Biomaterials. 2022;286:121611.

35. Boos MA, Lamandé SR, Stok KS. Multiscale Strain Transfer in Cartilage. Frontiers in Cell and Developmental Biology. 2022;10:795522.

36. Hashemi-Afzal F, Fallahi H, Bagheri F, Collins MN, Eslaminejad MB, Seitz H. Advancements in hydrogel design for articular cartilage regeneration: A comprehensive review. Bioact Mater. 2025;43:1–31.

37. Perez RA, Mestres G. Role of pore size and morphology in musculo-skeletal tissue regeneration. Mater Sci Eng C Mater Biol Appl. 2016;61:922–39.

38. Zhou L, Wang Z, Chen D, Lin J, Li W, Guo S, et al. An injectable and photocurable methacrylate-silk fibroin hydrogel loaded with bFGF for spinal cord regeneration. Materials & Design. 2022;217:110670.

39. Jian G, Li D, Ying Q, Chen X, Zhai Q, Wang S, et al. Dual Photo-Enhanced Interpenetrating Network Hydrogel with Biophysical and Biochemical Signals for Infected Bone Defect Healing. Adv Healthc Mater. 2023;12(25):e2300469.

40. Stepien A, Juszczak L, Koronowicz A, Such A, Kowalski G, Synkiewicz-Musialska B, et al. Effect of Hyaluronic Acid Content on Functional Properties, Antioxidant Activity, and In Vitro Digestion of Food-Grade Furcellaran Hydrogels and Emulgels. Materials. 2025;18(24):5581.

41. Menezes R, Vincent R, Osorno L, Hu P, Arinzeh TL. Biomaterials and tissue engineering approaches using glycosaminoglycans for tissue repair: Lessons learned from the native extracellular matrix. Acta Biomater. 2023;163:210–27.

42. Dovedytis M, Liu ZJ, Bartlett S. Hyaluronic acid and its biomedical applications: A review. Engineered Regeneration. 2020;1:102–13.

43. Pawcenis D, Koperska MA, Milczarek JM, Łojewski T, Łojewska J. Size exclusion chromatography for analyses of fibroin in silk: optimization of sampling and separation conditions. Applied Physics A. 2014;114(2):301–8.

44. Hong H, Lee OJ, Lee YJ, Lee JS, Ajiteru O, Lee H, et al. Cytocompatibility of Modified Silk Fibroin with Glycidyl Methacrylate for Tissue Engineering and Biomedical Applications. Biomolecules. 2020;11(1).

45. Kudva AK, Luyten FP, Patterson J. Initiating human articular chondrocyte re-differentiation in a 3D system after 2D expansion. J Mater Sci Mater Med. 2017;28(10):156.

46. Yao H, Tan R, Zhang Y, Wang C, Wan Y, Min Q. Multi-crosslinked hydrogels composed of hyaluronic acid, silk fibroin and chitosan nanoparticles with capacity of bioactive molecule delivery and degradation tolerance for cartilage tissue engineering. Eur Polym J. 2024;220:113449.

47. Li Y, Chen X, Zhou Z, Fang B, Chen Z, Huang Y, et al. Berberine oleanolic acid complex salt grafted hyaluronic acid/silk fibroin (BOA-g-HA/SF) composite scaffold promotes cartilage tissue regeneration under IL-1beta caused stress. Int J Biol Macromol. 2023;250:126104.

48. Ziadlou R, Rotman S, Teuschl A, Salzer E, Barbero A, Martin I, et al. Optimization of hyaluronic acid-tyramine/silk-fibroin composite hydrogels for cartilage tissue engineering and delivery of anti-inflammatory and anabolic drugs. Mater Sci Eng C Mater Biol Appl. 2021;120:111701.

49. Bandyopadhyay A, Mandal BB, Bhardwaj N. 3D bioprinting of photo-crosslinkable silk methacrylate (SilMA)-polyethylene glycol diacrylate (PEGDA) bioink for cartilage tissue engineering. J Biomed Mater Res A. 2022;110(4):884–98.

50. Kabir W, Di Bella C, Choong PFM, O’Connell CD. Assessment of Native Human Articular Cartilage: A Biomechanical Protocol. Cartilage. 2021;13(2_suppl):427S–37S.

51. Wan J, Jiang J, Yu X, Zhou J, Wang Y, Fu S, et al. Injectable biomimetic hydrogel based on modified chitosan and silk fibroin with decellularized cartilage extracellular matrix for cartilage repair and regeneration. Int J Biol Macromol. 2025;298:140058.

